# What underlies exceptional memory function in older age? No evidence for aging-specific relationships to hippocampal atrophy and retrieval activity

**DOI:** 10.1101/2024.04.12.589183

**Authors:** Anders M. Fjell, Markus H. Sneve, Inge K Amlien, Håkon Grydeland, Athanasia M. Mowinckel, Didac Vidal-Piñeiro, Øystein Sørensen, Kristine B. Walhovd

**Affiliations:** Center for Lifespan Changes in Brain and Cognition, Department of Psychology, University of Oslo, 0373 Oslo, Norway; Center for Computational Radiology and Artificial Intelligence, Department of Radiology and Nuclear Medicine, Oslo University Hospital, 0424 Oslo, Norway

**Keywords:** aging, episodic memory, hippocampus, magnetic resonance imaging, SuperAgers, brain activity, atrophy

## Abstract

Some older adults show superior memory performance compared to same-age peers, even performing on par with young participants. These are often referred to as *SuperAgers*. It is not known whether their superior memory function is caused by special features of their brains in aging, or whether superior memory has the same brain foundation throughout adult life. To address this, we measured hippocampal volume and atrophy, microstructural integrity by diffusion tensor imaging, and activity during an episodic memory encoding and retrieval task, in 277 cognitively healthy adults (age 20.1-81.5 years at baseline, mean 49.2 years). For quantification of hippocampal atrophy, all participants had repeated MRIs, from two to seven examinations, covering a mean of 9.3 years between first and last scan (2.5-17.3 years). 15.7% of the participants above 60 years had episodic memory scores above the mean of the young and middle-aged participants and were classified as SuperAgers. We found that superior memory in older adults was associated with higher retrieval activity in the anterior hippocampus and less hippocampal atrophy. However, there were no significant age-interactions, suggesting that the relationships reflected stable correlates of superior memory function. Although SuperAgers had superior memory compared to their same-age peers, they still performed worse than the best-performing young participants. Further, age-memory performance curves across the full age-range were similar for participants with superior memory performance compared to those with normal and low performance. These trajectories were based on cross-sectional data, but do not indicate preserved memory among the superior functioning older adults. In conclusion, the current results confirm that aspects of hippocampal structure and function are related to superior memory across age, without evidence to suggest that SuperAgers have special features compared to their younger counterparts.

## Introduction

Some older adults show superior memory function compared to same-age peers, and even perform on par with much younger persons (Powell, Lam, et al., 2023). These very high-performing older adults are sometimes referred to as *SuperAgers* (Harrison, Weintraub, Mesulam, & Rogalski, 2012). The aim of the present study was to explore hippocampal contributions to superior memory function in aging by investigating volume, rates of atrophy, microstructural integrity, and encoding– and retrieval related activity, which may all play a role in episodic memory function in aging (Nyberg & Pudas, 2019). In particular, we aimed to test whether there are exceptional features of the hippocampus in aging that can help explain the very high performance of SuperAgers, or whether the neural correlates of superior memory are similar across adult life.

A yet unanswered question is whether superior memory performance in aging is due to resilience to expected age-related brain decline (Nyberg, Lovden, Riklund, Lindenberger, & Backman, 2012), or rather reflects life-long high function. The latter explanation is in line with research showing high correlations between cognitive function in early and later life, that individual differences in brain and cognition appear very stable over the adult lifespan, and that differences in change rate appear to be a relatively weak source of individual differences among older adults (Karama et al., 2014; Vidal-Pineiro et al., 2021; Walhovd, Lovden, & Fjell, 2023). Successful aging may thus have a foundation in early life cognitive function which then is continuously shaped in response to changing environments (Nyberg & Pudas, 2019; Walhovd, Howell, Ritchie, Staff, & Cotman, 2019; Walhovd et al., 2023). However, some studies have also found high-functioning older adults to show less memory decline over time (Gefen et al., 2014; Harrison, Maass, Baker, & Jagust, 2018; Josefsson, de Luna, Pudas, Nilsson, & Nyberg, 2012), although with some variation (Dekhtyar et al., 2017). Hence, superior memory performance in older age may represent a mix of life-long high function and relatively less decline.

Few longitudinal brain imaging studies have addressed the topic of superior memory function directly, but cross-sectional studies have reported larger volumes of hippocampal and other brain structures when older high-performers are compared to their normal performing age-peers (Dekhtyar et al., 2017; Harrison et al., 2018; Harrison et al., 2012; Sun et al., 2016). Such findings may reflect stable individual differences and do not necessarily imply brain maintenance (Dekhtyar et al., 2017; Harrison et al., 2018), a question which requires longitudinal data to address. Less whole-brain cortical volume loss in high-performers was reported in one longitudinal study (Cook et al., 2017), and memory decline in aging is related to macro– and microstructural brain changes, especially of the hippocampus (Fjell et al., 2014; Langnes et al., 2020). Hippocampal volume loss typically accelerates in older adults (Fjell, Westlye, et al., 2013; Vinke et al., 2018), even in participants with very low risk of Alzheimer’s Disease (AD) (Fjell, Westlye, et al., 2013), with a lifespan trajectory mimicking that seen for episodic memory. Similarly, integrity of the hippocampus, as measured by diffusion tensor imaging (DTI), is reduced across the adult lifespan (Langnes et al., 2020). These hippocampal changes are not only benign, with more atrophy and integrity reductions being related to steeper decline of episodic memory, although the associations are usually modest (Fjell, McEvoy, et al., 2013; Gorbach et al., 2020; Gorbach et al., 2017).

Measures of brain activity during episodic memory tasks could explain part of the remaining variance in memory performance not accounted for by hippocampal structure and atrophy. Maintained memory function has been related to preservation of functional memory networks including the hippocampus (Duzel, Schutze, Yonelinas, & Heinze, 2011; Pudas et al., 2013; Vidal-Pineiro et al., 2019), although others have argued that compensatory brain activity can support memory function in aging (Davis, Dennis, Daselaar, Fleck, & Cabeza, 2008). Hippocampal activity measured during execution of episodic memory tasks may thus contribute to explain superior memory performance in some older adults.

## Materials and methods

### Sample

The participants were recruited from ongoing studies coordinated by the Center for Lifespan Changes in Brain and Cognition (LCBC) at the Department of Psychology, University of Oslo, Norway. The final sample consisted of all available participants February 5^th^, 2024 who were 20 years or older, with valid cross-sectional task-fMRI recordings during an associative encoding and retrieval task, diffusion tensor imaging (DTI) scans, and longitudinal T1-weighted MRIs for quantification of atrophy. The final sample consisted of 277 well-screened cognitively healthy participants (137 females, age 20.1-81.5 years at baseline, mean 49.2 years, SD = 17.7 years). 113 had two separate scanning sessions, 63 had 3, 65 had 4, 23 had 5, 7 had 6, and 6 had 7. The mean follow-up interval from baseline to the last scan was 9.3 years (2.5-17.3 years, SD = 4.2). The participants were compensated for their participation.

All participants gave written informed consent, and the Regional Ethical Committee of South Norway approved the study. At baseline, the participants reported no history of neurological or psychiatric disorders, chronic illness, premature birth, learning disabilities, or use of medicines known to affect nervous system functioning. They were further required to speak fluent Norwegian and have normal or corrected-to-normal hearing and vision, and were required to score ≥26 on the Mini Mental State Examination (Folstein, Folstein, & McHugh, 1975). Participants were tested on Vocabulary and Matrix Reasoning subtests of Wechsler’s Abbreviated Scale Intelligence Scale (WASI) (Wechsler, 1999), and all scored within the normal IQ range (>85). Participants were further excluded due to experimental and operator errors, low number of trials available for fMRI analysis (n = 24 from the total participant pool; <6 trials per condition of interest) and extreme head movement (n = 1; >1.5 mm mean movement), reducing the final sample to the above mentioned 277 participants. The sample partially overlaps with the samples used in Sneve et al. (Sneve et al., 2015), Vidal-Piñero et al. (Vidal-Pineiro et al., 2017), and (Langnes et al., 2019), which addressed different research questions.

### Experimental design – fMRI tasks

Participants were scanned using BOLD fMRI during an associative encoding task and an unannounced subsequent memory test after ≈ 90 minutes. After the encoding, participants were taken out of the scanner, and not given any specific instructions about what to do during the retention interval. Importantly, they were not informed that they would be tested for the encoded material. The fMRI task is described in detail elsewhere (Sneve et al., 2015). Briefly, the encoding and the retrieval tasks consisted of two and four runs, respectively, of 50 trials each. All runs started and ended with an 11s baseline period, which was also presented once in the middle of each run. The stimulus material consisted of 300 black and white line drawings depicting everyday objects and items. During encoding, the participants went through 100 trials and performed simple evaluations. A trial had the following structure: a female voice asked either “Can you eat it?” or “Can you lift it?” Both questions were asked equally often and were pseudorandomly mixed across the different objects. One second after question onset, a black and white line drawing of an object was presented on the screen along with response indicators, and participants were instructed to produce yes/no– responses. Button response was counterbalanced across participants. There were no correct or incorrect responses. The object remained on the screen for 2 s, when it was replaced by a central fixation cross that remained throughout the intertrial interval (ITI; 1-7s exponential distribution over four discrete intervals).

During the surprise memory test, 200 line drawings of objects were presented, of which an equal number was old or new. A test trial started with the pseudorandom presentation of an object and the auditory presented question (Question 1) “Have you seen this item before?”. Each object stayed on the screen for 2 seconds, and participants were instructed to respond “Yes” or “No”. If the participant responded “Yes”, a new question followed (Question 2): “Can you remember what you were asked to do with the item?” A “Yes”-response to this question led to a final two-alternative forced choice question (Question 3): “Were you asked to eat it or lift it?” Here, the participant indicated either “Eat” or “Lift”. The design efficiency was tentatively optimized to ensure sufficient complexity in the recorded time series (http://surfer.nmr.mgh.harvard.edu/optseq/).

### Analysis of behavioral data

Multiple behavior variables were extracted and used in the analyses. Number of correct recognized items (regardless of source), correct rejections, recognition misses, false alarms, and recollections were entered into a principal component analysis to extract a main component of memory performance which was used as the memory measure of interest in all analyses. This component explained 53.9% of the variance, with recollection responses showing the highest loading (.96).

### MRI scanning

At baseline, imaging was performed at a Siemens Skyra 3T MRI with a 20-channel head-neck coil at Oslo University Hospital, Rikshospitalet. For functional imaging the parameters were equivalent across all runs: 43 slices (transversal, no gap) were measured using T2* weighted BOLD EPI (TR=2390ms; TE=30ms; flip angle=90°; voxel size=3×3×3mm; FOV=224×224; interleaved acquisition; GRAPPA=2). Each encoding run produced 131 volumes while the number of volumes per retrieval run was dependent on participants’ responses (mean 207 volumes). Three dummy volumes were collected at the start of each fMRI run to avoid T1 saturation effects in the analyzed data. Additionally, a standard double-echo gradient-echo field map sequence was acquired for distortion correction of the EPI images. Visual stimuli were presented in the scanner environment with a 32-inch InroomViewing Device monitor while participants responded using the ResponseGrip device (both NordicNeuroLab, Norway). Auditory stimuli were presented to the participants’ headphones through the scanner intercom.

Anatomical T1-weighted MPRAGE images consisting of 176 sagittally oriented slices were obtained using a turbo field echo pulse sequence (TR = 2300 msec, TE = 2.98 msec, flip angle = 8°, voxel size = 1 × 1 × 1 mm, FOV= 256 × 256 mm). For DTI, a single-shot twice-refocused spin-echo echo planar imaging (EPI) with 64 directions: TR = 9300 ms, TE = 87 ms, b-value = 1000 s/mm^2^, voxel size = 2.0 × 2.0 × 2.0 mm, slice spacing = 2.6 mm, FOV = 256, matrix size = 128 × 130 × 70, 1 non-diffusion-weighted (b = 0) image. Another non-diffusion-weighted (b=0) image was acquired with the reverse phase encoding for distortion correction.

Longitudinal data included anatomical T1-weighted scans only. At follow-up, some participants were scanned on a Siemens 3T Prisma (208 sagittally oriented slices using a turbo field echo pulse sequence: TR = 2400 ms; TE = 2.22 ms; flip angle = 8°; voxel size = 0.8 × 0.8 × 0.8 mm). We have previously shown that the correlations between hippocampal volume across the Skyra and the Prisma are high (r ≥ .85) (Langnes et al., 2020). In the present study, we regressed out the effect of scanner on hippocampal volume and did all statistical analyses on the residuals. As a test of the procedure, we correlated hippocampal offset volume calculated only from Skyra scans with offset volumes including both Skyra and Prisma scans at the latest timepoints. The correlation was r = .99. Hence, the approach of regressing out scanner change from the longitudinal anatomical data seems valid.

### MRI preprocessing and analysis

Volumetric segmentation of the hippocampus from the T1-weighted scans was performed with FreeSurfer 7.1 (https://surfer.nmr.mgh.harvard.edu/) (Fischl et al., 2004a, 2002). DTI scans were processed with FMRIB’s Diffusion Toolbox (fsl.fmrib.ox.ac.uk/fsl/fslwiki) (Jenkinson, Beckmann, Behrens, Woolrich, & Smith, 2012; Smith et al., 2004). From the B0 images with reversed phase-encode blips, we estimated the susceptibility-induced off-resonance field using a method similar to what is described in (Andersson et al., 2003) as implemented in FSL (Smith et al., 2004). We then applied the estimate of the susceptibility induced off-resonance field with the eddy tool (Andersson and Sotiropoulos, 2016), which was also used to correct eddy-current induced distortions and participant head movement, align all images to the first image in the series and rotate the bvecs in accordance with the image alignments performed in the previous steps (Jenkinson et al., 2002; Leemans and Jones, 2009). After estimating the diffusion ellipsoid properties, i.e., the three eigenvalues defining the length of the longest, middle, and shortest axes of the ellipsoid, we computed mean diffusivity (MD), defined as the average of the three eigenvalues, as implemented in the FSL tool *dtifit*. The processed MD maps were co-registered to the segmented T1-weighted images to extract average MD values in the hippocampus. Specifically, for each participant, all DTI voxels for which more than 50% of the underlying anatomical voxels were labeled as hippocampus by FreeSurfer were considered representations of the hippocampus.

fMRI data was initially corrected for susceptibility distortions, motion and slice timing corrected, smoothed (5mm FWHM) in volume space and high-pass filtered (at 0.01Hz) using FSL (http://fsl.fmrib.ox.ac.uk/fsl/fslwiki). Next, FMRIB’s ICA-based Xnoiseifier (FIX, v1.06) (Salimi-Khorshidi et al., 2014) was used to auto-classify noise components and remove them from the fMRI data. The classifier was trained on a task-specific dataset in which task fMRI data from 36 participants had been manually classified into signal and noise components (age span in training set: 7-80; fMRI acquisition parameters identical to the current study). Motion confounds (24 parameters) were regressed out of the data as a part of the FIX routines. Transformation matrices between functional-native, structural-native and FreeSurfer average space were computed to delineate hippocampal structures and bring them to the functional-native space.

The preprocessed fMRI data was entered in a first-level general linear model (GLM) with FSFAST (https://surfer.nmr.mgh.harvard.edu/fswiki/FsFast) for each encoding and retrieval run, consisting of several conditions/regressors modeled as events with onsets and durations corresponding to the trial events during encoding and retrieval and convolved with a two-gamma canonical hemodynamic response function (HRF). At retrieval, each “old” trial (test item presented during encoding, n = 100) was assigned to a condition based on the participant’s response at test. Two conditions of interest were modeled both at encoding and at retrieval. 1) The source memory encoding condition consisted of items that were later correctly recognized with correct source memory (Yes response to test Questions 1 and 2 and correct response to Question 3). 2) The miss condition consisted of items presented during encoding which were not recognized during test (incorrect No response to test Question 1). In addition, several regressors were included to account for BOLD variance associated with task aspects not included in any investigated contrast. During both encoding and retrieval, an item memory condition was included that consisted of items that were correctly recognized but for which the participant had no source memory (Yes response to Question 1 and No response to test Question 2 or incorrect response to Question 3) as well as a fourth regressor that modeled trials in which the participant did not produce any response to the first question. For the retrieval runs, four additional regressors were included to model the response to the new items (i.e. correct rejections and false alarms) and to model the second and third test questions (Questions 2 and 3). Temporal autocorrelations [AR(1)] in the residuals were corrected using a prewhitening approach. For memory analyses, a contrast of interest consisting of source – miss memory conditions was computed for each participant.

### Hippocampal segmentation for fMRI

Since encoding and retrieval are known to activate anterior and posterior sections of the hippocampus to various degrees, we divided the FreeSurfer segmented hippocampus in an anterior and a posterior part. Moving anteriorly through the coronal planes of an MNI-resampled human brain, y = –21 corresponds to the appearance of the uncus of the parahippocampal gyrus. In line with recommendations for long-axis segmentation of the hippocampus in human neuroimaging (Poppenk, Evensmoen, Moscovitch, & Nadel, 2013), we labeled hippocampal voxels at or anterior to this landmark as anterior HC while voxels posterior to the uncal apex were labeled as posterior HC. Specifically, for each participant, all functional voxels for which more than 50% of the underlying anatomical voxels were labeled as hippocampus by FreeSurfer were considered functional representations of the hippocampus. While keeping the data in native subject space, we next established hippocampal voxels’ locations relative to MNI y = –21 by calculating the inverse of the MNI-transformation parameters for a given subject’s brain and projecting the back-transformed coronal plane corresponding to MNI y = –21 to functional native space. All reported activity– and connectivity measures thus represent averages from hippocampal sub-regions established in native space

### Definition of memory performance groups

Memory performance was indexed by the principal component of all the response variables from the retrieval-fMRI task (see above). We used two different approaches to define memory performance groups. First, we followed the most common approach used in previous research on SuperAgers, which is to classify high-performing older adults as participants above 60 years with memory scores above the average of the participants in the age-range 20 to 60 years (Powell, Page, Close, Sachdev, & Brodaty, 2023). Note that there is no uniform definition of SuperAgers, and a review reported that frequency of SuperAgers varies from 2.9% to 43.4% as a function of differences in definition (Powell, Lam, et al., 2023). It has been argued that a start age of 60 years, rather than 80 years, which was used in the original work on SuperAgers (Harrison et al., 2012), is problematic when the aim is to make meaningful statements about resilience and resistance to age-decline (Rogalski, 2019). Still, as cognitive and brain aging is ongoing well before 60 years (Fjell et al., 2014), and there is a high correlation between early and late-life cognitive function and brain health (Karama et al., 2014; Walhovd et al., 2023), it is likely that relevant brain correlates of superior memory function in higher age can be identified from 60 years. Unless there are unique processes that can only been seen after 80 years, so that the high cognitive function of SuperAgers has different brain causes or correlates before this age, a start age of 60 years should be a reasonable choice considering the vast reductions in sample size by using a cut-off of 80 years.

Hence, we calculated mean memory performance in participants 20 to 60 years and used this to split the sample in two older (≥ 60 years) and two younger/middle-aged groups (20 to 60 years). High-functioning older adults (> 60 years, n = 15, 4 females) were defined as those with higher than average memory scores for young/ middle-aged adults, and normal-functioning older adults were defined as the rest of the older sample (n = 80, 48 females). Similarly, the young sample was split in two based on memory score above the mean (n = 97, 63 females) or below the mean (n = 85, 58 females). Descriptive statistics for the groups are provided in Table 1.

**Table 1.** Descriptive group statistics. Working mem is the principal component (z-score) of out-of-scanner working memory tests, i.e. digit span forward, digit span backward and letter memory. Recall mem is the principal component (z-score) of out-of-scanner memory tests, i.e. CVLT total learning, 30 min free recall and RCFT 20 min recall. P-values for the post hoc ANOVA tests were adjusted by Tukey HSD to keep a 95% family-wise confidence level.

|  | Older adults |  | Younger adults |  | Group comparisons |  |  |
| --- | --- | --- | --- | --- | --- | --- | --- |
|  | Group 1:<br>High | Group 2:<br>Normal | Group 3:<br>High | Group 4:<br>Normal | 1 - 2<br>*p* = | 3 - 4<br>*p* = | ANOVA<br>*p* = |
| n | 15 | 80 | 97 | 85 |  |  |  |
| Age | 67.7 | 69.1 | 37.7 | 40.12 | .97 | .42 |  |
| Sex (f/m) | 4/11 | 48/32 | 63/34 | 58/27 | .07 | .97 | 0.19 |
| Working mem (z) | -0.36 | -0.38 | 0.42 | -0.06 | .99 | .002 | 1.1e<sup>-7</sup> |
| Recall mem (z) | 0.14 | -0.68 | 0.42 | 0.15 | .004 | .15 | 1.4e<sup>-14</sup> |

Defining groups based on an age cut-off does not respect the continuous age-related variation in memory scores, making it difficult to address age-related variance. Therefore, we also created three groups based on age-corrected memory performance. We regressed out age from the memory score, and defined participants scoring ≥1 SD above the age-corrected mean as high performers, 1 SD ≤ below as low performers, and the rest as normal performers. This allowed us to test SuperPerformers across the age-range, instead of restricting this label to those above 60 years. By this approach, we could test if the brain correlates of superior memory had commonalities across adulthood, or whether there are unique aspects to superior memory function in older age.

### Statistical analyses

Analyses were run in R (https://www.r-project.org) using Rstudio (www.rstudio.com) IDE, except for an ANOVA which was run in IBM SPSS 29.0.2.0. For each participant, we calculated hippocampal slope (“atrophy”) and offset (“volume”) by fitting a linear model with hippocampal volume as dependent and time since baseline as independent variable. We then corrected hippocampal offset variables (volume) for estimated intracranial volume by running a linear regression and saving the residuals. All statistical analyses were done on the ICV-corrected hippocampal volumes.

Hippocampal activity was measured separately for anterior vs. posterior hippocampus and encoding vs retrieval. We therefore ran a repeated-measures ANOVA with phase (two levels: encoding, retrieval) × longitudinal axis (two levels: anterior, posterior) as within-subject factors and age group (two levels: older, young/ middle-aged) and memory performance group (2 levels: high, normal) as between-subject factors to disentangle the relationships to memory from the different fMRI variables. Sex and age were included as covariates. In a previous study using an overlapping sample we found only anterior hippocampal retrieval activity to be nominally related to memory performance (Langnes et al., 2019). However, that sample was much larger with a wider age-range (6.8-80.8 years) and used a different metric for memory performance, so it was necessary to test the relationship within this study.

Generalized Additive Models (GAM) using the package “mgcv” (Wood, 2006) were used to derive age-functions and to test age-interactions. Group means in memory and the neuroimaging markers were compared using Tukey’s Honestly Significant Difference (Tukey HSD) method yielding adjusted p-values to control the familywise error rate (95%). We first used the age-dependent four-groups memory performance classification (SuperAgers), and then repeated the main analyses using the three-group age-independent performance classification (SuperPerformers). Finally, we ran a GAM with memory score as continuous variable and all the hippocampal variables as predictors, with age, sex, and six variables indexing movement during the fMRI scanning as covariates.

## 3.1 Results

### Descriptive group statistics

There were no significant group differences in age or sex within each age category. The high performing older group had a higher score than the normal performing older group on a PCA generated from two independent out-of-scanner memory tests (the California Verbal Learning Test learning and 30 min recall conditions (Delis, Kramer, Kaplan, & Ober, 2000) and the Rey-Osterrieth Complex Figure Test 20 min recall condition (Meyers & Meyers, 1995)), demonstrating that superior memory performance was not restricted to the in-scanner memory test used for group classification. There were no differences between the two older groups on a PCA of two out-of-scanner working memory tests (principal component of digit span forward, backward, and the letter memory test) (Krogsrud et al., 2021).

### Distribution of memory scores and age-relationships

The Shapiro-Wilk normality test suggested that the memory scores did not deviate from a normal distribution (W = 0.99, p = 0.34) (Figure 1). This suggests that all participants are drawn from the same underlying distribution of memory scores, and that there are no distinct subgroups within the overall sample which exhibit different patterns of performance on the memory test.

**Figure 1.**
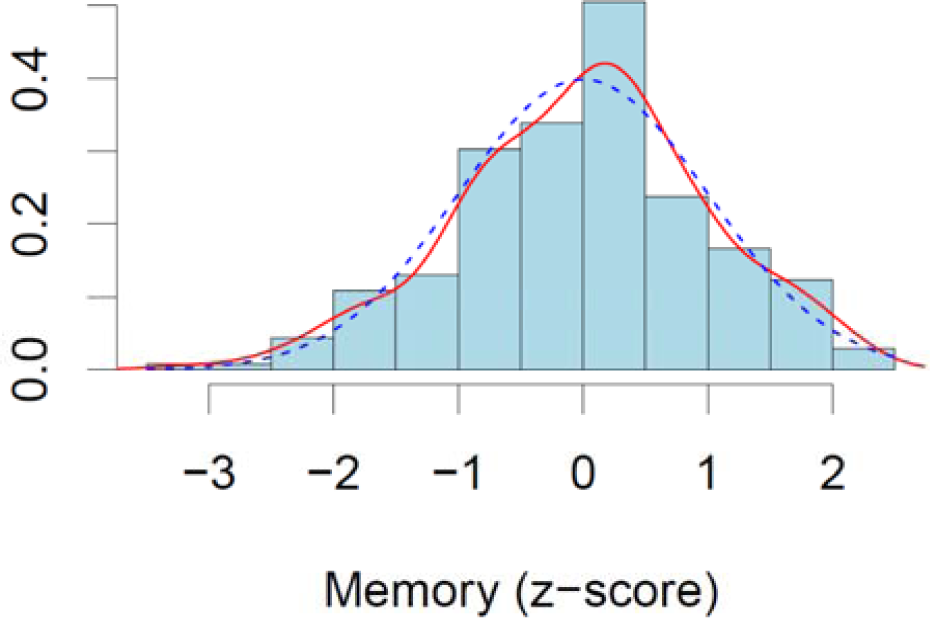
Density plot of unadjusted memory scores. The red curve shows the density distribution of the data, and the blue dotted line shown the normal distribution. Memory performance (z-scores) is on the x-axis, density on the y-axis.

Memory was linearly and negatively related to age (Estimate = –0.023 SD change pr year, SE = 0.003, t = –7.88, p < 7.87e^-14^), with a correlation of r = –.42. The age-trajectory is shown in Figure 2 (top left panel), and the individual scores are displayed with color codes according to age-dependent (left panel) or age-independent (right panel) performance group.

**Figure 2.**
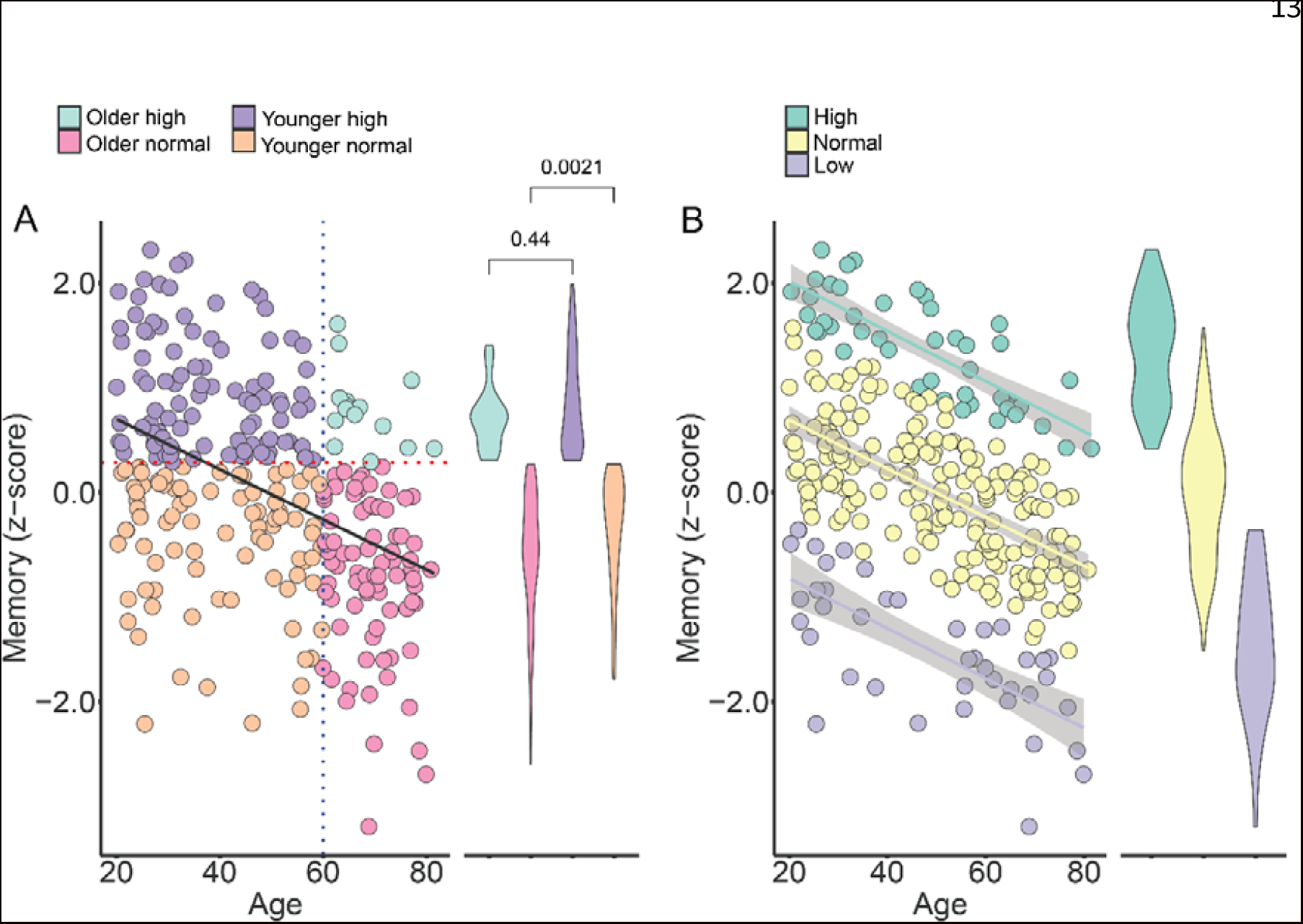
Memory across age. Scatterplots showing individual memory scores across age, with mean and distribution of scores for each performance group illustrated by the violin plots next to each scatterplot. Panel A: Dots are color-coded according to age group (vertical blue dotted line: above vs. below 60 years) and memory performance (horizontal red dotted line: above vs. below the mean score of the young/ middle-aged group). P-values show for comparisons between the two high performing groups, and between the two normal performing groups. Group 1 (cyan): Older high performing; Group 2 (light pink): Older normal performing; Group 3 (dark purple): Young/ middle-age high performing. Group 4 (beige): Young/ middle-age normal performing. Panel B: Color coding according to age-adjusted memory performance. Group 1 (green): high function; Group 2 (yellow): normal function; Group 3 (light purple): low function. The shaded areas around the lines depict 95% CI.

### Age-relationships of hippocampal volume, atrophy, microstructure, and activity

Except for memory-related brain activity, all hippocampal features were significantly related to age, with steeper relationships from mid-life. Scatterplots are shown in Figure 3 and numeric results in Table 2. There was a correlation between larger hippocampal volume and less atrophy (r = .21, p < .001).

**Figure 3.**
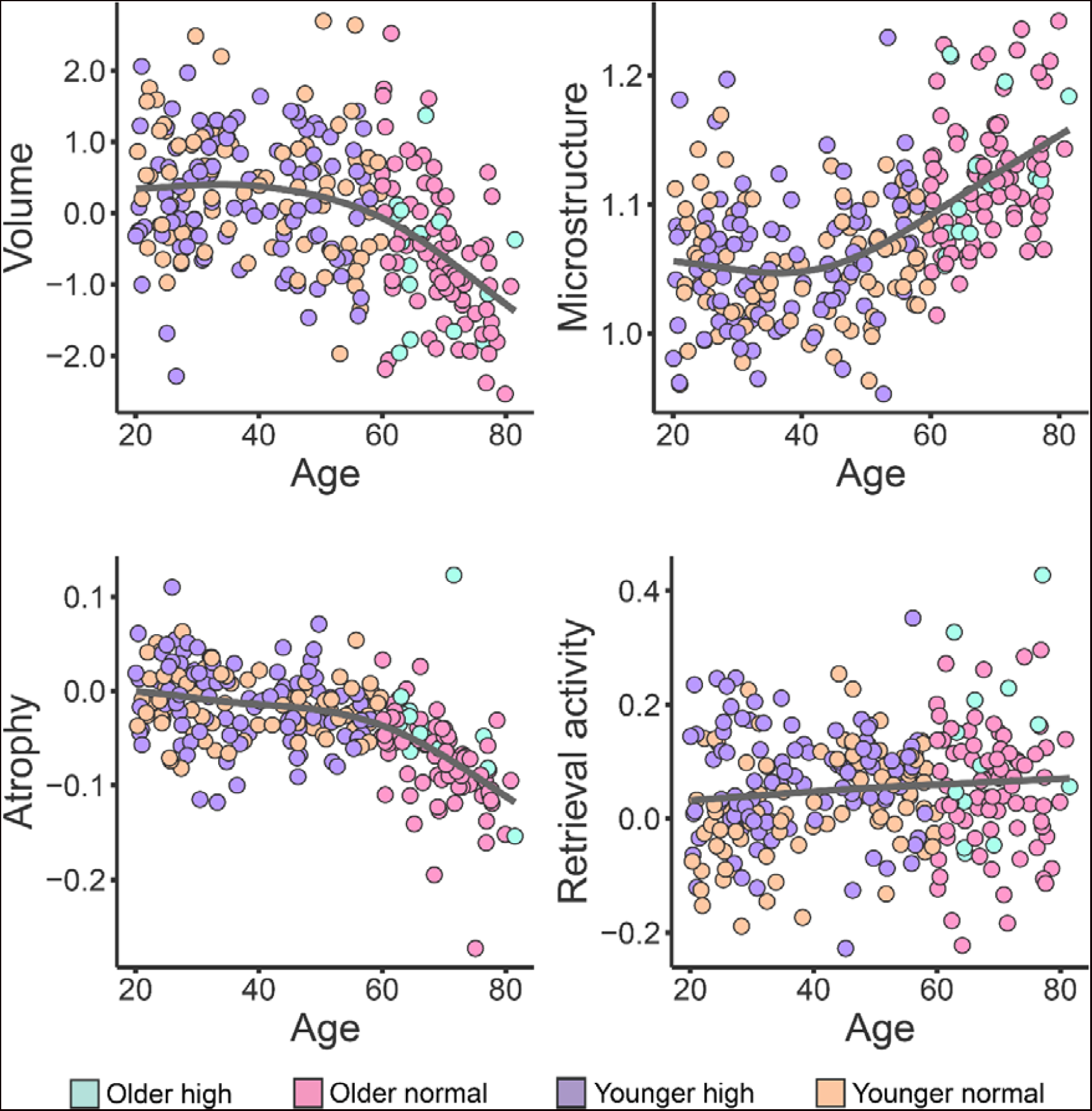
Age-relationships of hippocampal features. Scatters are color-coded by age-dependent memory performance group identity. Retrieval activity was measured in the anterior hippocampus as the contrast between successfully recollected items and forgotten items (miss). Group 1 (cyan): Older high performing; Group 2 (light pink): Older normal performing; Group 3 (purple): Young/ middle-age high performing. Group 4 (beige): Young/ middle-age normal performing. The unit for volume is z-score, for microstructure mm^2^/s ×1000, for atrophy annual change in z-score.

**Table 2.** Age-relationships of hippocampus features. Results of repeated GAMs with each hippocampal feature as dependent and s(Age) as independent variable. Edf: Effective degrees of freedom (expressing degree of non-linearity). P-values not adjusted for multiple comparisons.

| Hippocampal feature | edf | F | R<sup>2</sup> | P< |
| --- | --- | --- | --- | --- |
| Volume (offset) | 2.70 | 24.03 | 0.23 | 2e<sup>-16</sup> |
| Atrophy (slope) | 3.17 | 40.26 | 0.37 | $2e^{-16}$ |
| Microstructure (MD) | 2.95 | 36.47 | 0.33 | $2e^{-16}$ |
| Encoding-anterior | 1.00 | 0.00 | 0.00 | 0.96 |
| Encoding-posterior | 1.00 | 3.20 | 0.01 | 0.07 |
| Retrieval-anterior | 1.18 | 2.14 | 0.01 | 0.1 |
| Retrieval-posterior | 4.10 | 1.84 | 0.03 | 0.1 |

### Age-dependent memory performance groups

The older high performing group (SuperAgers) had comparable memory to the younger high performing group (older high: z = 0.80 [SE = 0.09] vs. younger high: z = 0.96 [SE = 0.06], Cohen’s D = 0.30 [CI: 0.25-0.84]). The older normal group had significantly poorer memory than the younger normal group (Cohen’s D = 0.47 [CI: 0.15-0.77]), see violin plots in Figure 2.

Group-wise comparisons of the hippocampal features are shown in Figure 4. Both older groups had lower ICV-corrected hippocampal volume and higher MD than the young groups (all p’s < .05, adjusted). The normal performing older group had more atrophy than all other groups, while there were no other differences in atrophy between any of the other groups (older normal vs older high: Cohen’s D = 0.74, CI: .17-1.30; older normal vs young high: Cohen’s D = 1.35, CI: 1.02-1.67; older normal vs young normal: Cohen’s D = 1.48, CI: 1.13-1.82). As can be seen in Figure 3, one high performing older participant had very low rate of atrophy, and one normal performing older participant had very high rate of atrophy. When removing these from the analyses, atrophy in the older low performers was not significantly different from the older high performers, while the older high performers now showed significantly higher rate of atrophy than both young groups (both p’s < .02). Whether these atrophy rates reflect real brain change or noise, is difficult to decide, but we decided to include these participants in further analyses. Still, as these two participants greatly affected the results, the results of the hippocampal atrophy analyses must be interpreted with this in mind. No other significant group differences were observed.

**Figure 4.**
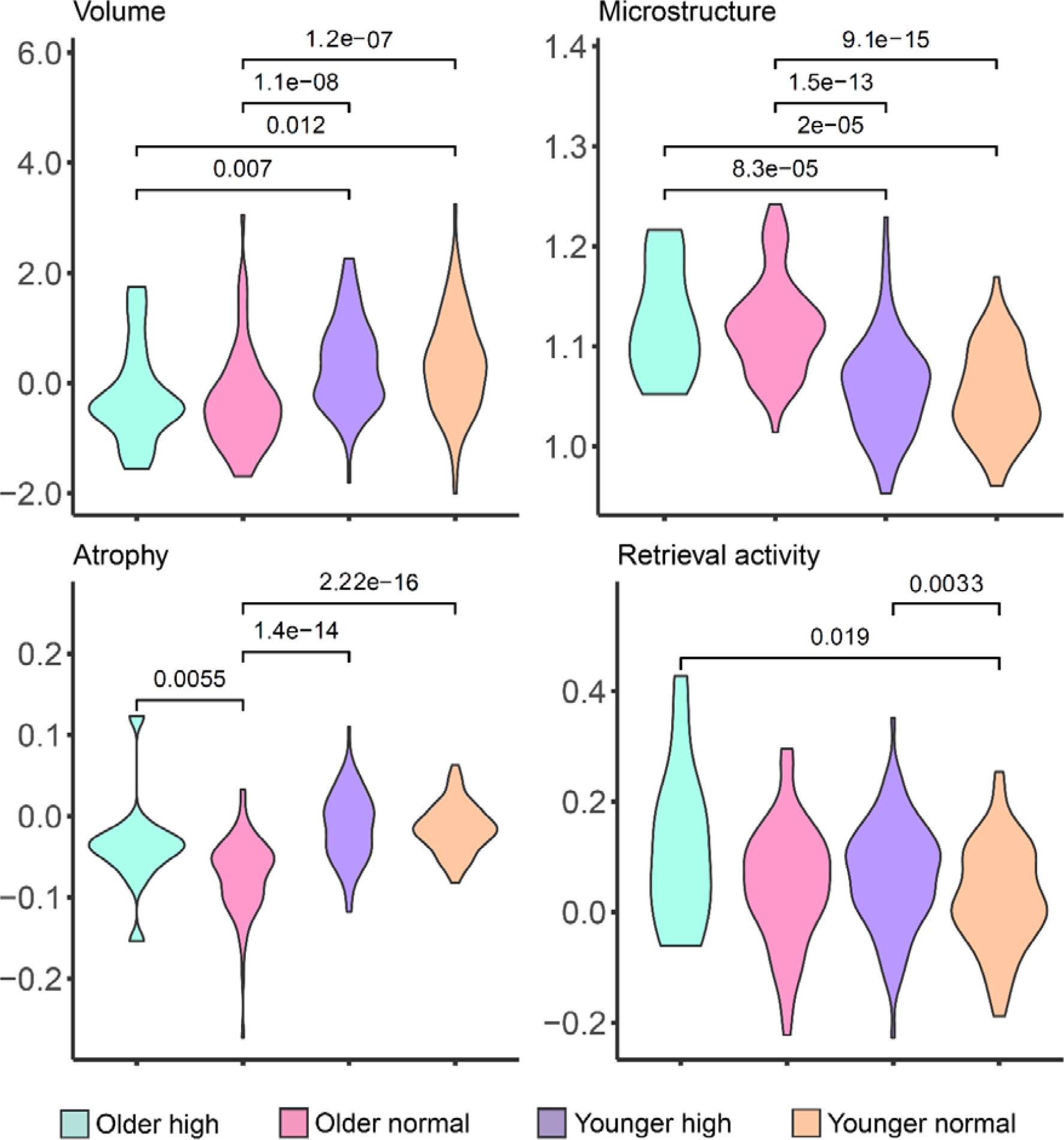
Violin plots illustrating the pairwise group comparisons for each hippocampal feature. P-values are corrected according to Tukey HSD. The groups are from left to right: Group 1 (cyan): Older high performing; Group 2 (light pink): Older normal performing; Group 3 (purple): Young/ middle-age high performing. Group 4 (beige): Young/ middle-age normal performing.

For brain activity, we ran a repeated-measures ANOVA with phase (two levels: encoding, retrieval) × longitudinal axis (two levels: anterior, posterior) as within-subject factors and age group (two levels: older, younger/ middle-aged) and memory performance group (2 levels: high, normal) as between-subject factors. Sex and age were included as covariates. There was a significant memory group × phase × long axis interaction (F = 5.91, p < .016), caused by larger effect of memory group on activity during retrieval than encoding, and this effect was larger in anterior than posterior hippocampus (see Figure 5). Post-hoc comparisons showed higher anterior retrieval-related activity in the older high vs. younger normal group (p < .05, adjusted), as well as nominally significant higher activity than for the older normal group, which did not survive Tukey HSD (adjusted p = 0.08). Younger high performers showed higher activity than younger normal performers. There were no significant interactions between memory group and age group. Full ANOVA-results are presented in SI Table 1.

**Figure 5.**
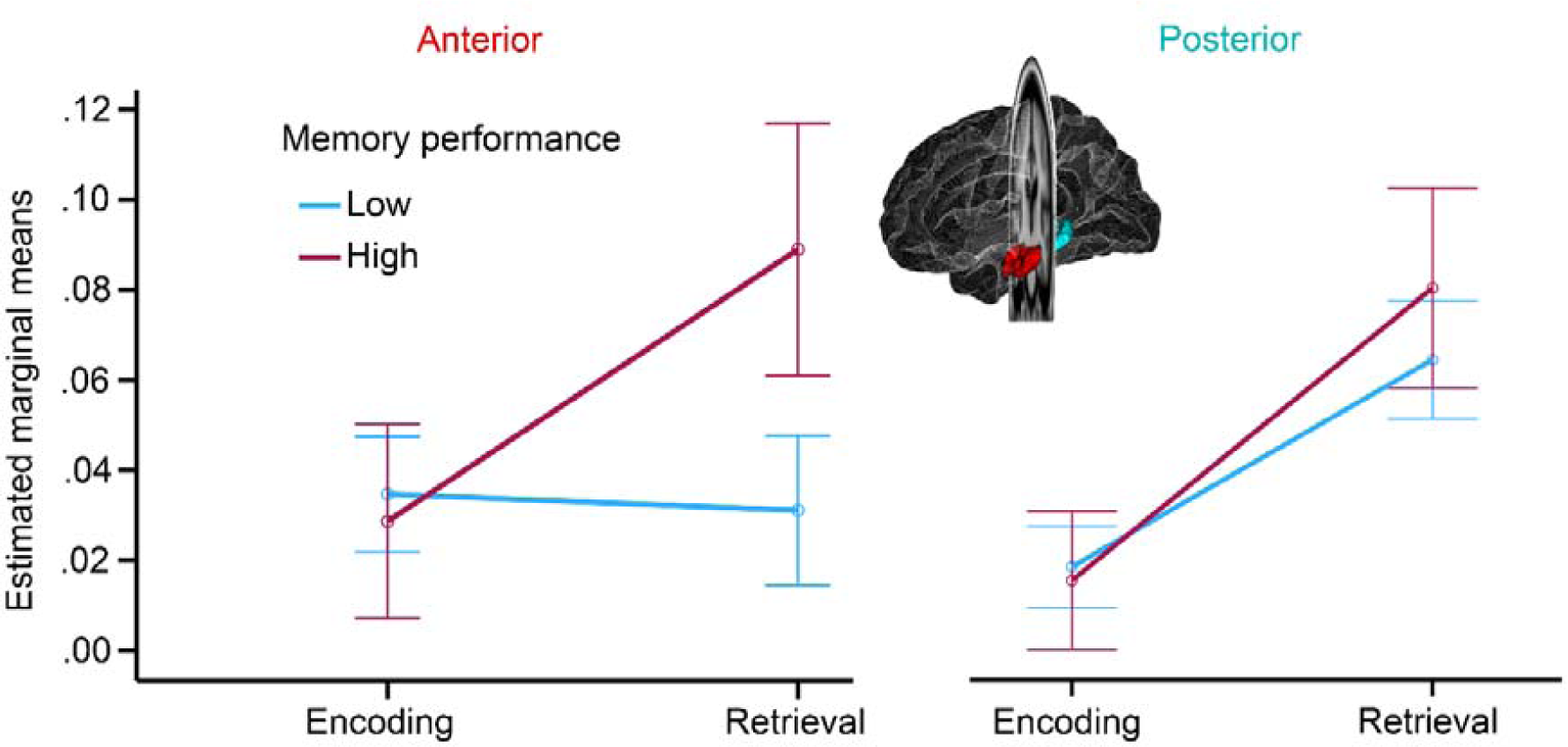
ANOVA of interactions between memory performance and memory-related fMRI activity. The figure illustrates that memory-related activity differences between participants with high vs. low memory performance were largest for the retrieval phase in the anterior hippocampus. Error bars depict 95% CI.

In summary, the results suggest that hippocampal atrophy and retrieval related activity are associated with superior memory function in aging, with the caveats regarding atrophy rate mentioned above. However, the analyses were not optimized for testing whether these associations were unique to aging, or whether they rather reflect stable, age-independent relationships. To address this, we re-analyzed the data using a group division based on age-independent memory performance.

### Age-independent memory performance groups

After regressing memory score on age, the residuals were used to assign the participants to groups of high, normal, and low memory performance. As can be seen in Figure 2 (Panel B), the age-trajectories for each performance group were close to parallel. The corresponding correlations between memory performance and age in each group was r = –.80 (high performers), r = –.66 (average performers), and r = –.72 (low performers). Main effects and age-interactions for group membership were tested by GAMs with each of the hippocampal features in turn as outcome variable (see Figure 6 for scatterplots of age-trajectories with observations colored by performance group). There was a significant effect of memory performance group on hippocampal atrophy (estimate = 0.014, t = 2.29, p = .023), due to more atrophy in the low than high performing group. Performance group was also significantly related to retrieval activity (estimate = 0.06, t = 4.02, p = 7.34e^-5^), with both the normal and low performing group showing lower activity than the high performing group. No significant age-interactions were detected. There were no main effects or age-interactions for hippocampal volume or microstructure. Running the analyses using memory performance as a continuous variable gave the same main results. All GAMs were run with both linear and tensor interactions to attempt to identify age-interactions, but no significant effects were seen. The age-invariant effects of memory performance group on atrophy and retrieval activity are illustrated in Figure 7.

**Figure 6.**
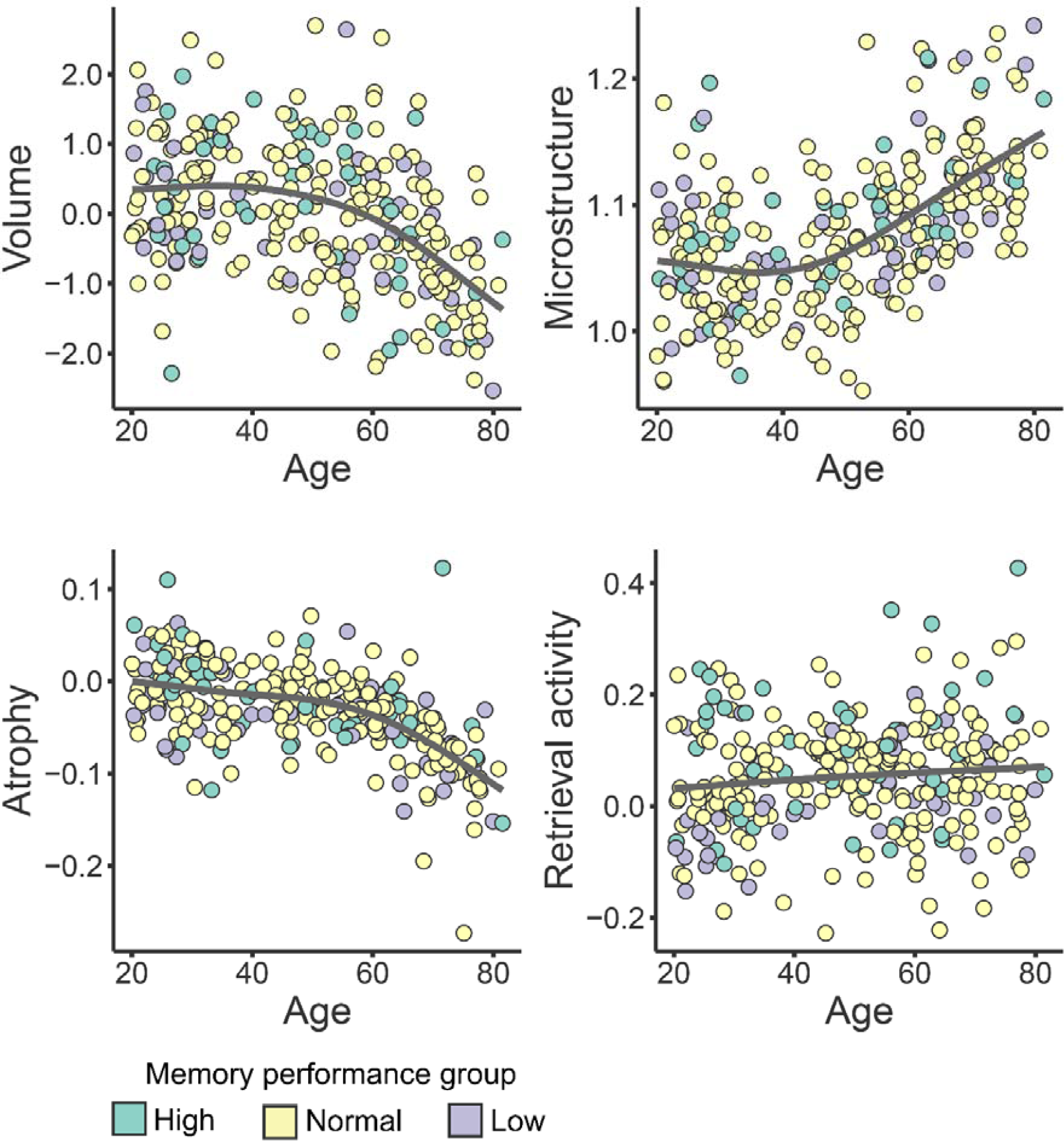
The same age-trajectories as in Figure 3, with scatters color-coded according to age-independent memory performance. The unit for volume is z-score, for microstructure mm^2^/s ×1000, for atrophy annual change in z-score. Group 1 (green): high function; Group 2 (yellow): normal function; Group 3 (dark purple): low function).

**Figure 7.**
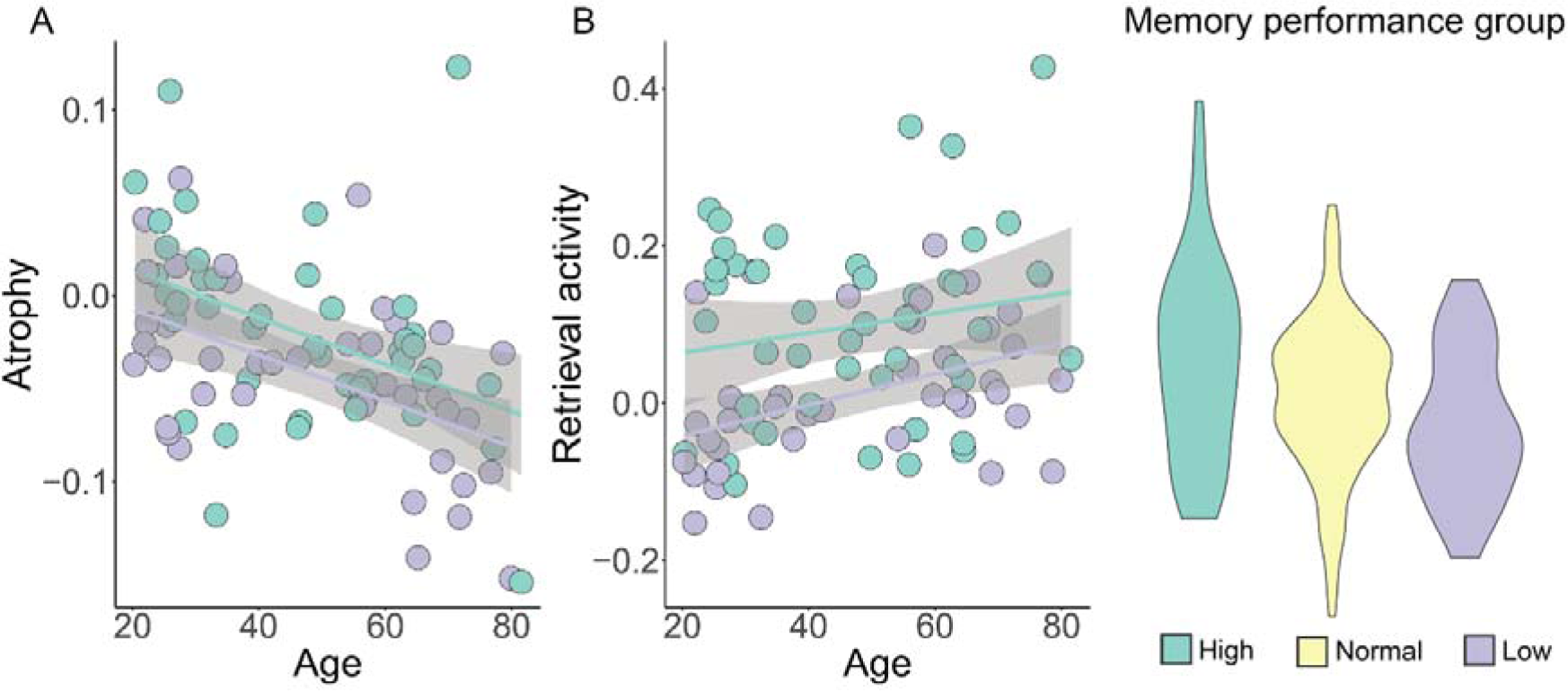
Age-trajectories for the high (green) and low (dark purple) memory performers for atrophy (left panel: A) and retrieval activity (middle panel: B). Violin plots showing the values and distribution of retrieval activity for the three age-independent memory performance groups, with the high perming group showing higher activity than the two other groups. The shaded areas around the lines depict 95% CI.

Finally, the GAM with all hippocampal features as simultaneous predictors of memory score showed that hippocampal atrophy (p = .045) and anterior hippocampus retrieval activity (p = 5.38e^-5^) were unique predictors. R^2^ (adjusted) for the total model was 0.233. We also re-ran the model, including average mean diffusion in the TBSS-derived skeleton as an additional predictor, which had minor effects on the results, reducing the p-value for atrophy slightly (.031), while retrieval activity was still highly significant (p = 4.46 e^-5^) and R^2^ (adjusted) = 0.237.

## Discussion

The results suggest that the same features of the hippocampus were related to superior memory across adult life, not being specific to aging. Superior memory performance showed stable associations to memory-related retrieval activity and hippocampal atrophy, and SuperAgers did not seem to represent a distinct sub-group with different brain correlates compared to super-performers in other age-groups. Results were generally in agreement whether participants were divided in groups based on age and memory performance (SuperAgers), age-independent memory performance (SuperPerformers), or memory was analyzed as a continuous variable. Hence, analyzing population representative participants across wide age-ranges and to study normal variation using continuous variables instead of creating groups based on various definitions may be preferable. Implications of the results are discussed below.

### Who have superior memory in aging?

Older adults with superior memory almost certainly had superior memory also when young. We know that there is a high correlation between cognitive function early and late in life (Arbuckle, Maag, Pushkar, & Chaikelson, 1998; Gow et al., 2011; Rönnlund, Sundström, & Nilsson, 2015; Walhovd et al., 2023), with age-invariant brain structural correlates (Walhovd et al., 2016). Studies focusing on episodic memory find that many SuperAgers are not able to maintain cognitive function, and almost none transform from normal agers to SuperAgers (Dekhtyar et al., 2017; Habib, Nyberg, & Nilsson, 2007). Although this cannot be directly tested in the present data of cross-sectional memory tests, it is notable that the relationship between memory and age was similar across the three age-independent performance groups. This shows that the difference between the SuperAgers and their younger counterparts of SuperPerformers is comparable to the differences between the other groups of younger and older participants. From this perspective, being a SuperAger mainly requires a high starting point, while slower age-expected decline may yield some additional benefit. This is in accordance with a recent study reporting larger cortical area across life in participants with high cognitive functions, but also some evidence of less change over time (Walhovd et al., 2022).

An influential model holds that maintenance of brain structure and function in late life is a primary condition for successful memory aging (Nyberg et al., 2012). One study showed that although SuperAgers experienced significant cortical atrophy over 18 month, volume loss was less than in age-matched controls (Cook et al., 2017). Here we see indications of the same phenomenon when comparing hippocampal atrophy between high and normal performing older adults. However, it needs to be stressed that one SuperAger with much less than expected hippocampal change and one older adult with normal memory score and more than expected hippocampal atrophy appeared to highly influence the association. When plotted across the full age-range, the atrophy-memory association turned out not to be age-specific. Thus, the results do not give strong evidence for low rate of atrophy being an important aspect of superior memory performance. Still, the number of SuperAgers was low, and several previous studies have shown significant, although modest, correlations between rate of hippocampal atrophy and change in episodic memory function in aging (Fjell, McEvoy, et al., 2013; Gorbach et al., 2020; Gorbach et al., 2017). Thus, low rate of hippocampal atrophy may be one correlate of relatively better preserved memory function, although it explains only a minor portion of the inter-individual differences in cognitively normal participants at any point in life.

Further evidence for brain maintenance comes from studies reporting that preservation of functional memory networks including the hippocampus is associated with better memory function in aging (Duzel et al., 2011; Pudas et al., 2013). In a previous study using an overlapping sample, Vidal-Piñeiro et al. found that frontal brain function during encoding resembling activity of younger participants was a primary characteristic of higher memory function in aging (Vidal-Pineiro et al., 2019). In the present study, the strongest association with high episodic memory function was found with anterior hippocampal retrieval activity. Memory-related activity was the only hippocampal feature not related to age, and the relationship with memory appeared as a stable association across the age-span. As illustrated in Figure 7, the age-curves of high vs. low performers are parallel. This suggests that at least some of the brain basis of superior memory in aging is also associated with superior memory in young and middle age. Concurrent longitudinal data on memory function spanning several years is needed to draw stronger conclusions.

Based on the current results, there may be limited knowledge to gain by focusing on SuperAgers instead of utilizing the full specter of memory scores across the lifespan. This may yield more statistical power and allow testing age-effects on brain-cognition associations with better precision. Our tentative division of participants +/− 1SD from the age-corrected mean fits reasonably well with a previous study using a data-driven pattern-mixture model to divide older participants in groups of maintainers (18%), decliners (13%) and average decliners (68%) based on change in performance on episodic memory tasks (Josefsson et al., 2012). Still, we acknowledge that there can be economic and practical arguments in longitudinal studies for following select samples of participants.

Several limitations of the present work must be highlighted. First, as memory performance was based on cross-sectional data, results do not directly inform about memory change and resistance to age-decline. Second, we focused on the hippocampus as this is a key region for memory function, with established importance in aging (Nyberg & Pudas, 2019). However, many other brain regions contribute to high memory function, and it is possible that some of these would show age-variant relationships to cognition. Finally, the employed definition of SuperAgers yielded a small sample with limited power to detect modest effects, such as the expected associations with cross-sectional volumetric measures and hippocampal microstructure (Klinedinst et al., 2023).

## Conclusion

Superior memory function in aging is related to features of hippocampal activity and atrophy in age-invariant ways. Future studies should assess changes in memory function with longitudinal designs coupled to multi-modal neuroimaging to disentangle stable factors from brain changes in older age cognitive function.

## Acknowledgements

This work was supported by the Department of Psychology, University of Oslo (to K.B.W., A.M.F.), the Norwegian Research Council (to K.B.W., A.M.F.) and the project has received funding from the European Research Council’s Starting/ Consolidator Grant schemes under grant agreements 283634, 725025 (to A.M.F.) and 313440 (to K.B.W.).

## Notes

### Competing Interest Statement

The authors have declared no competing interest.

